# Bleomycin induced EBV-specific gastric epithelial cell death is enhanced by LMP2A

**DOI:** 10.1101/2024.01.23.576907

**Authors:** Aung Phyo Wai, Afifa Fatimah Azzahra Ahmad Wadi, Yuxin Liu, Tumurgan Zolzaya, Shunpei Okada, Hisashi Iizasa, Hironori Yoshiyama

**Affiliations:** Department of Microbiology, Faculty of Medicine, Shimane University, Shimane, Japan

**Keywords:** EBV, EBV-associated epithelial cancer, lytic infection, LMP2, Bleomycin, DDR, apoptosis

## Abstract

Epstein-Barr virus (EBV) infects more than 90% of the global population, leading to EBV-related cancers, such as EBV-associated gastric cancer and nasopharyngeal carcinoma. Despite the rarity of p53 mutations in tumor cells, there is a lack of specific therapies targeting EBV-associated gastric cancers. Our study demonstrated that EBV-infected gastric epithelial cells exhibited high sensitivity to the bleomycin family, encompassing bleomycin and zeocin, compared to uninfected cells. Zeocin treatment induces a DNA damage response (DDR) and promotes the expression of BZLF1, triggering the expression of EBV lytic genes. This cascade leads to apoptosis via a p53 dependent pathway, coinciding with the activation of caspase-3. Cells infected with the BZLF1-deficient virus could not induce apoptosis by zeocin treatment, and the cells maintained a tight latent infection but initiated a robust DDR. Conversely, neither lytic infection nor DDR was induced by zeocin treatment in cells infected with LMP2A-deficient virus. These findings indicate that LMP2A augments DDR in EBV-infected gastric epithelial cells, facilitating the transition from latent to lytic infection, and consequently inducing cell death. Bleomycin derivatives are expected to serve as fundamental components in the development of specific anticancer drugs targeting EBV-positive epithelial cancer cells.

**IMPORTANCE:** Because of its ubiquitous infection in the human population, the number of patients suffering from EBV-related diseases, including malignant tumors and severe active infections, is substantial. Previous research has reported the potential of drugs such as HDAC inhibitors and poly ADP-ribose polymerase inhibitors to induce cell death in EBV-infected cells. However, a clinical therapeutic agent specifically reducing EBV-infected cells has not yet been available. In this study, we report that bleomycin, or its derivative, which has been used for over half a century, induces apoptotic cell death in EBV-infected cells by triggering lytic viral replication associated with DNA damage response. Particularly, we discovered that the viral latent gene LMP2A unexpectedly enhances bleomycin-induced EBV-infected epithelial cell death by promoting DDR.

---

Epstein-Barr virus (EBV), a human gamma-herpesvirus, persistently infects >90% of the adult population and is strongly associated with the development of several human cancers, such as Burkitt’s lymphoma, EBV-infected natural killer/T lymphomas, Hodgkin’s lymphoma, nasopharyngeal carcinoma, and EBV-associated gastric cancer (EBVaGC) (1). EBV-related malignancies develop in latently infected cells. The interplay between various viral and host genes, along with the involvement of various cellular pathways, leads to cancer (2).

EBV exhibits a biphasic infection within the host cell, latent and lytic form of infection. During latent infection, a limited set of latent viral genes is expressed. Inhibition of apoptosis, immune evasion, and induction of cell proliferation also occur during latent and lytic EBV infection (3). Reactivation from latent to lytic infection is initiated by the immediate early protein *Bam*HI Z fragment leftward open reading frame 1 (BZLF1), which activates a cascade of viral genes to express >80 genes for viral production (4). Lytic EBV infection causes host cell apoptosis after a large amount of virus is released from the host cells (1).

As EBV must replicate its genomic DNA within the host cell nucleus, the virus has evolved mechanisms to use the host DNA damage response (DDR) for successful infection and replication. DDR induces phosphorylation of ataxia-telangiectasia mutated (ATM), enhances expression of histone H2AX phosphorylated on Ser193 (γH2AX), and mediates DNA damage checkpoint protein 1(5–7). These proteins induce p53 and activate caspase-3, -6, and -7, resulting in the induction of apoptosis (8). When EBV infection causes DNA damage, ATM is activated as a part of the DDR machinery of the host cell. ATM activation leads to phosphorylation and activation of p53, which positively regulates EBV lytic replication (9–13). ATM activation also enhances expression of the EBV lytic BZLF1 gene, promoting the switch from latent to lytic replication (13). However, the detailed mechanisms and specific triggers governing the transition from the different latency types to the lytic phase of EBV infection are not well known.

EBVaGC, a subtype of gastric cancer, accounts for approximately 9% of all gastric cancers and is associated with a better prognosis than other EBV-associated malignancies. Infrequent mutation in p53 gene compared to other types of gastric cancers is a characteristic of EBVaGC (14–16). EBVaGC also responds well to chemotherapeutic agents, often inducing DDR and lytic infections via activation of ATM and p53 (17–19). EBVaGC shows latent type I infection and expresses latent membrane protein 2A (LMP2A), Epstein-Barr nuclear antigen 1, *Bam*HI A fragment right transcript microRNAs, and EBV-encoded small RNAs (20).

LMP2A is a membrane-anchored protein with an immunoreceptor tyrosine-based activation motif in its intracellular N-terminal region. LMP2A mimics the signaling of the precursor B cell receptor (pre-BCR), which also contains immunoreceptor tyrosine-based activation motif, and triggers the activation of mitogen-activated protein kinase, phosphoinositide 3-kinase, and Akt via Src family tyrosine kinases such as Lyn and Syk (21). LMP2A expressed on EBV-infected B lymphocytes hijacks the pre-BCR signal and maintains latent infection by preventing EBV from entering a lytic infection (22). LMP2A expressed on epithelial cells is involved in tumorigenesis by acquiring anoikis resistance and decreasing phosphatase and tensin homolog expression (23, 24). However, few reports have described the role of LMP2A in the EBV life cycle during epithelial infection.

We noticed that bleomycin strongly induced apoptosis in latent EBV-infected gastric epithelial cells. Bleomycin is an antitumor antibiotic derived from *Streptomyces* (25). Bleomycin causes double-stranded DNA breaks, a type of DDR, by inducing oxidative stress (26–28). Zeocin is an analog of bleomycin and is more stable than bleomycin. However, zeocin has been experimentally used to induce double-stranded DNA breaks because of its toxicity. We elucidated the molecular mechanism by which the bleomycin family preferentially injures EBV-positive gastric epithelial cells and discuss the potential application of the bleomycin family in oncolytic therapy for EBVaGC.

## MATERIALS AND METHODS

### Cell culture and drugs

AGS cells from human stomach cancer and Daudi cells from Burkitt’s lymphoma were obtained from the American Type Culture Collection (Manassas, VA, USA). MKN28 cells were obtained from the National Institute of Biomedical Innovation, Health, and Nutrition (JCRB Cell Bank). Immortalized fetal gastric epithelial GES1 cells were obtained from the Beijing Institute for Cancer Research. We also used EBV-negative Daudi cells (29). AGS cells were infected with a recombinant Akata EBV strain, whose viral thymidine kinase BXLF1 gene was disrupted by inserting the neomycin resistance gene and enhanced green fluorescent protein (eGFP) gene (eGFP-EBV) (30). Cells were cultured at 37℃ in a 5% CO_2_ using RPMI-1640 medium (Sigma-Aldrich, St. Louis, MO, USA) supplemented with 10% fetal bovine serum (Sigma-Aldrich), 100 units/mL of penicillin, and 100 μg/mL of streptomycin (Nacalai, Kyoto, Japan). To maintain EBV-positive GES1 (GES1-EBV) cell and EBV-positive MKN28 (MKN28-EBV) cell, 250 and 560 μg/mL of G418 (Promega, Madison, WI, USA) was added to the culture medium, respectively. To maintain EBV-positive AGS (AGS-EBV) cell, LMP2A knockout (KO) EBV-infected AGS (AGS-EBV LMP2A KO) cell, BZLF1 KO EBV-infected AGS (AGS-EBV BZLF1 KO) cell, 500, 500, and 560 μg/mL of G418 was used, respectively (31,32). Zeocin solution was purchased from Invitrogen (Waltham, MA, USA. USA). Bleomycin sulfate powder (Tokyo Chemical Industry, Tokyo, Japan) was dissolved in water.

### Cell cytotoxicity assay

One hundred microliters of EBV-positive or -negative gastric epithelial cells suspended at 5 × 10^4^ cells/mL were seeded into 96-well plates and cultured overnight at 37 °C in a 5% CO_2_ incubator. Then, the cells were treated with zeocin and incubated for 36 h at 37 °C in a 5% CO_2_ incubator. Ten microliters of CCK-8 solution (2-[2-methoxy-4-nitrophenyl]-3-[4- nitrophenyl]-5-[2, 4-disulfophenyl]-2H-tetrazolium) (DOJINDO, Kumamoto, Japan) were added to each well. The plates were incubated at 37 °C for 3 h, and the absorbance at 450 nm was measured using a microplate reader (DTX880, Beckman coulter, Brea, CA, USA). The half-maximal inhibitory concentration (IC_50_) was calculated using CCK-8 data and GraphPad Prism version 9 (GraphPad Software, Boston, MA, USA).

### Flow cytometry

To seed 1.5 × 10^5^ cells per well, 2.5 mL of AGS-EBV cells or AGS cells suspended at 6 × 10^4^ cells/mL were poured into each well of a six-well plate and cultured at 37 °C in a 5% CO_2_ incubator. Then, cells were treated with 200 μg/mL of zeocin or 10 μg/mL of bleomycin. Apoptotic cells were detected using a Dead Cell Apoptosis Kit (Thermo Fisher Scientific) and a Cytoflex flow cytometer (Beckman Coulter). 7-amino-actinomycin D was used as a marker for dead cells. Annexin V is used as an early marker of apoptosis.

### Immunoblot analysis

EBV-positive or -negative gastric epithelial cells were treated with 200 μg/mL of zeocin or 10 μg/mL of bleomycin. RIPA buffer (Thermo Fisher) was added along with a protease inhibitor (cOmplete mini, Sigma-Aldrich) and a phosphatase inhibitor (PhosStop, Sigma-Aldrich) to lyse the cells. Protein samples weighing 10 μg were electrophoresed using 7.5, 10, 12, or 15% SDS-polyacrylamid gels, then transferred to PVDF membranes (Merk Millipore, Brea, CA, USA). PhosphoBlocker Blocking Reagent (Cell Biolabs, San Diego, CA, USA), BlockAce (KAC, Amagasaki, Japan), or Tris-buffered saline with 0.01% Tween 20 and 5% bovine serum albumin was used to block the membranes. The membranes were then incubated with antibodies against γH2AX (20E3; Cell Signaling Technology (CST), Danvers, MA, USA), ATM (MAT3-4G10/8; Biolegend, San Diego, CA, USA), phospho-ATM (p-ATM) (Ser1981) (D25E5; CST), p53 (DO1; Active Motif, Carlsbad, CA, USA), phospho-p53 (Ser15) (16G8; CST), caspase-3 (W20054B; Biolegend), cleaved caspase-3 (Asp175) (5A1E; CST), EBV BZLF1 (BZ.1; Dako, Glostrup, Denmark), LMP2A (15F9; Abcam Cambridge, UK), glyceraldehyde triphosphate dehydrogenase (GAPDH) (EPR16891; Abcam), and β-actin (C4; Merk Millipore). After washing, membranes were incubated with horseradish peroxidase-conjugated anti-rabbit IgG (CST), horseradish peroxidase-conjugated anti-rat IgG (Beckman Coulter), or horseradish peroxidase-conjugated anti-mouse IgG (CST) antibodies. Specific signals were visualized using Immobilon Chemiluminescent reagent (Merck Millipore) and X-ray films. GAPDH or β-actin was used as an internal control.

### Viral titration assay

AGS-EBV cells were treated with or withoutzeocin for 48 h and centrifuged at 400 × *g* for 5 min at room temperature. The collected supernatant was sterilized using a 0.45-μm membrane filter, then concentrated at 16,000 × *g* for 90 min at 4 °C. After removing the supernatant, the pellet was resuspended with a new medium and kept at -80 °C until use. To measure the virus infectious titer, 10-fold serial dilutions of the viral supernatant were infected to Daudi (-) cells. FACS analysis was performed 48 h after EBV infection to detect the cells expressing eGFP (33). AGS-EBV cells treated with bleomycin or zeocin were examined.

### Small RNA interference (siRNA) treatment

AGS or AGS-EBV cells were suspended with 4 × 10^4^ cells/mL and seeded with 250 µL on each well of a 48-well plate for overnight (33). All siRNAs were obtained from Integrated DNA Technologies (IDT Coralville, IA, USA). TP53 siRNAs (2 nmol/L) (hs.Ri.TP53.13.1: sense, 5’-AACCUCUUGGUGAACCUUAGUACCT-3’ antisense, 5’- AGGUACUAAGGUUCACCAAGAGGUUGU-3’ or hs.Ri.TP53.13.2: sense, 5’-AGCAUCUUAUC CGAGUGGAAGGAAA-3, antisense, 5’- UUUCCUUCCACUCGGAUAAGAUGCUGA-3’ were transfected with RNAiMaX (Thermo Fisher Scientific) according to the manufacturer’s protocol. After 48 h, the efficacy of the siRNAs was confirmed using western blotting. siRNA-treated cells were cultured with or without zeocin, and their cytotoxicity was examined after 36 h.

### Reverse transcriptase-quantitative polymerase chain reaction

Total RNAs was extracted using the ISOGEN reagent (Nippon Gene, Tokyo, Japan), following the manufacturer’s instructions. Subsequently, complementary DNAs was synthesized using SuperScript III reverse transcriptase (Thermo Fisher Scientific) and random hexamers in accordance with the manufacturer’s protocol. EBV gene expression was determined by the following primers purchased from IDT: *BZLF1* (F: 5’- TCCGACTGGGTCGTGGTT-3’, R: 5’-GCTGCATAAGCTTGATAAGCATTC-3′), *BALF5* (F: 5’- TAGGGCCAGTCAAAGTTG-3’, R: 5’-ACCTGCGAAGACATAGAG-3’), *BLLF1* (F: 5’-GTCAG TACACCATCCAGAGCC-3’, R: 5’-TTGGTAGAGAGCCTTCGTATG-3’), and *GAPDH* (F: 5′-AA TCCCATCACCATCTTCCA-3’, R: 5’-TGGACTCCACGACGTACTCA-3’) (33). Reverse transcriptase-quantitative polymerase chain reaction was performed using SsoAdvanced Universal SYBR Green Supermix (Bio-Rad Laboratories). Amplified target genes were detected using the CFX Connect real-time PCR detection system (Bio-Rad) (40 cycles of 98 °C for 2 min, 95 °C for 10 s, and 60 °C for 30 s). The expression level of GAPDH was used as the internal standard.

### Quantitative PCR

Genomic DNA was isolated using a GeneElute mammalian genomic DNA miniprep kit (Sigma-Aldrich) according to the manufacturer’s protocol. Quantitative PCR was performed using primers (*BamH* I W F: 5’-CCCAACACTCCACCACACC-3’ R: 5’-TCTTAGGAGG CTGTCCGAGG-3’ and GAPDH gene F: 5’-TGTGCTCCCACTCCTGATTTC-3’, R: 5’-CCTAGTC CCAGGGCTTTGATT-3’) and double-quenched fluorescent DNA probes (*Bam*H I W: 5’-FAM/CA CACACTA/ZEN/CACACACCCACCCGTCTC/IBFQ-3’, GAPDH: 5’-FAM/CGGTCACAA/ZEN/ TCTCCACGC/IBFQ-3’), respectively. The GAPDH gene was used as an internal standard (34).

### LMP2A gene transfection

One microliter of AGS cells suspended at 5 × 10^5^ cells/mL were poured into each well of a six-well plate and cultured at 37 °C in a 5% CO_2_ incubator. Four micrograms of either pcDNA3.1 or pcDNA3.1-HA-LMP2A plasmid DNAs (provided by Prof. T. Kanda, Tohoku Medical and Pharmaceutical University) were transfected with Lipofectamine 2000 (Thermo Fisher Scientific) according to the manufacturer’s protocol. Forty-eight hours after transfection, cells were treated with 200 μg/mL of zeocin for 3 h and subjected with protein extraction followed by immunoblot analysis as described above.

### Statistical analysis

All data were analyzed using GraphPad Prism version 9. Data were analyzed for statistical significance using two-way analysis of variance multiple comparison tests. *P* <0.05 was considered statistically significant.

## RESULTS

### Zeocin induces preferential cell cytotoxicity to EBV-positive gastric cancer cells

The IC_50_ of zeocin on EBV-positive AGS (AGS-EBV) cells and EBV-negative AGS cells were 306 and 1,015 µg/mL, respectively (Fig. 1A, left). Similarly, the IC_50_ of zeocin on GES1-EBV cells and EBV-negative GES1 cells were 163 and 1,014 µg/mL, respectively (Fig. 1A, right). We also observed that bleomycin exhibited stronger cytotoxicity against AGS-EBV cells than against AGS cells (Fig. 2A). Treating cells with 200 µg/ml of zeocin induced apoptosis for AGS-EBV cells at 35.70%, but AGS cells were at 4.97% (*P* <0.001) (Fig. 1B). Treating cells with 100 µg/ml of zeocin also induced apoptosis for GES1-EBV cells at 27.27%, but for GES1 cells at 7.22% (*P* <0.001) (Supplementary Fig. 1A). Consistently, treatment of cells with 10 µg/ml of bleomycin induced apoptosis for AGS-EBV cells at 33.08% and for AGS cells at 7.83% (*P* <0.001) (Fig. 2B).

**FIG 1.**
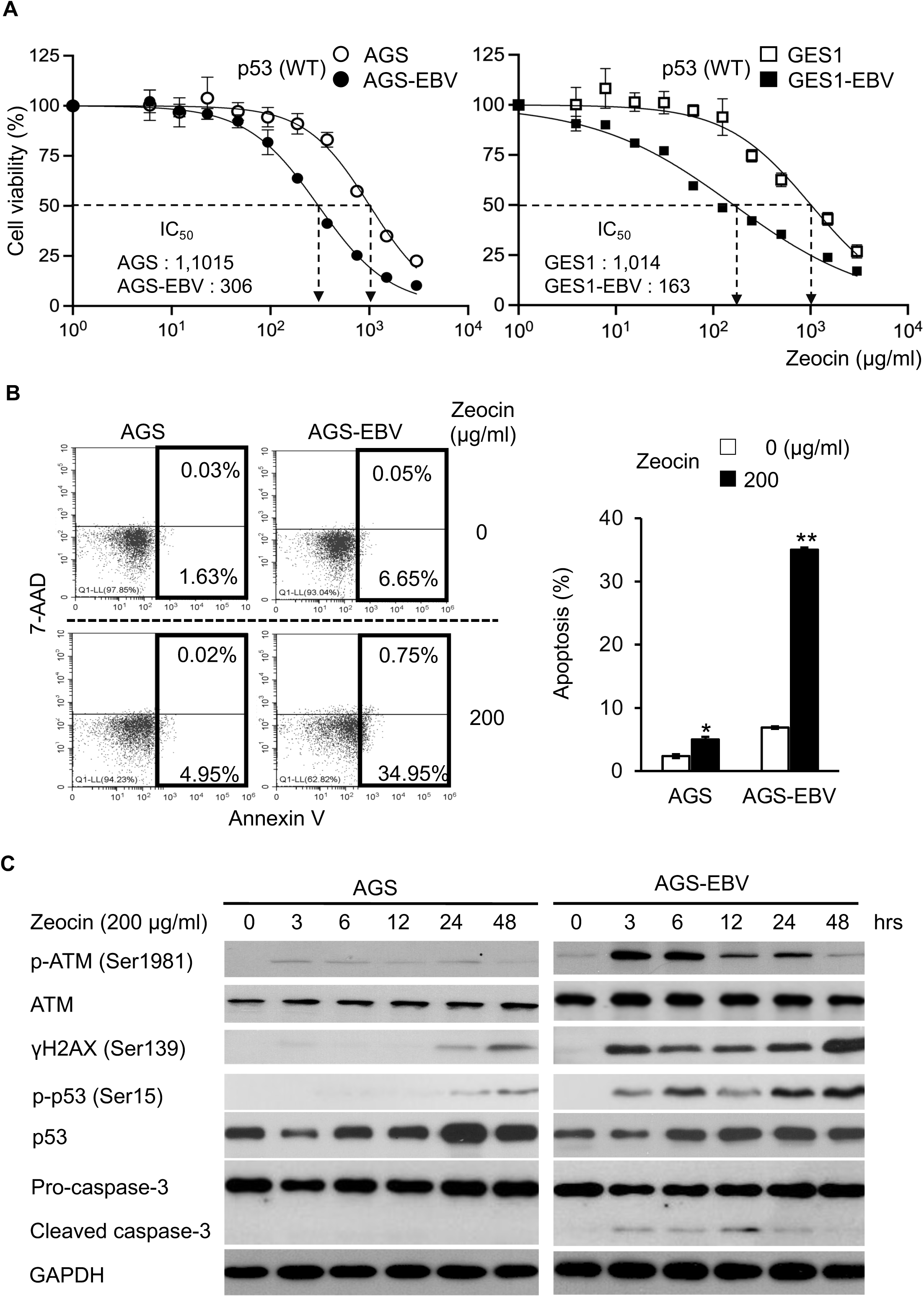
Zeocin induces stronger DDR in EBV-positive gastric cells than EBV-negative cells. **(A)** Cell viability in EBV-negative and -positive cells having normal p53 after zeocin treatment. Cell viability was compared with control DMSO treated cells. White circle: AGS cells, Black circle: AGS-EBV cells, White square: GES1 cells, Black square: GES1-EBV cells. **(B)** Apoptosis after zeocin treatment in AGS cells and AGS-EBV cells. Left: FACS analysis, Early apoptosis: 7-AAD^-^/Annexin V^+^, Late apoptosis: 7-AAD^+^/Annexin V^+^. Right: Apoptosis frequency after zeocin treatment. Apoptosis was the sum of early and late apoptosis. **(C)** Expression of DDR-related proteins in AGS cells and AGS-EBV cells after zeocin treatment. GAPDH was used as internal control. All experiments are representative of at least three independent experiments. _*_: *P* <0.05, _**_: *P* <0.01.

**FIG 2.**
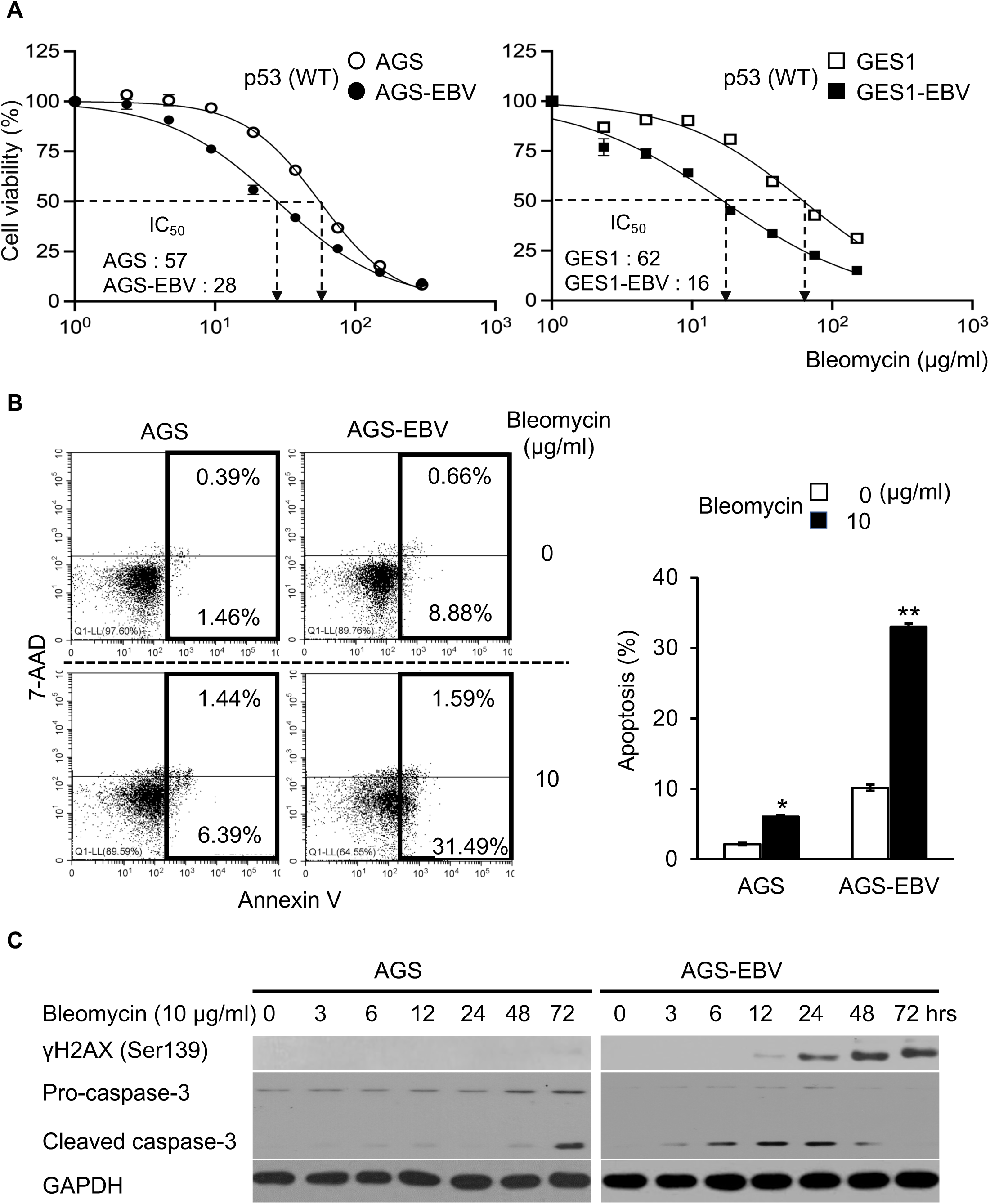
Bleomycin induces stronger DDR in EBV-positive gastric cells than EBV-negative cells. **(A)** Cell viability in EBV-negative and -positive cells having normal p53 after bleomycin treatment. Cell viability was compared with control DMSO treated cells. White circle: AGS cells, Black circle: AGS-EBV cells, White square: GES1 cells, Black square: GES1-EBV cells. **(B)** Apoptosis after bleomycin treatment in AGS cells and AGS-EBV cells. Left: FACS analysis, Early apoptosis: 7- AAD^-^/Annexin V^+^, Late apoptosis: 7-AAD^+^/Annexin V^+^. Right: Apoptosis frequency after bleomycin treatment. Apoptosis was the sum of early and late apoptosis. **(C)** Expression of DDR-related proteins in AGS and AGS-EBV cells after bleomycin treatment. GAPDH was used as internal control. All experiments are representative of at least three independent experiments. _*_: *P* <0.05, _**_: *P* <0.01.

The cytotoxicity of the bleomycin family is initiated by double-stranded DNA breaks (35, 36). Strong p-ATM signals were detected in AGS-EBV cells as early as 3 h after zeocin treatment (Fig. 1C). Moreover, the expression of γH2AX, which is a downstream factor of p-ATM that indicates DNA damage, was detected 3 h after zeocin treatment in AGS-EBV cells. However, expression of γH2AX could not detected in AGS cells until 24 h after zeocin treatment (Fig. 1C). Additionally, the expression of phospho-p53, another downstream molecule of p-ATM, increased 3 h after zeocin treatment in AGS-EBV cells (Fig. 1C). Expression of cleaved caspase-3, which is located downstream of phospho-p53 and induces apoptosis, was observed only in AGS-EBV cells treated with zeocin. Expression of cleaved caspase-3 was also observed in GES1-EBV cells treated with zeocin (Supplementary Fig. 1B).

Bleomycin also induced both γH2AX and cleaved caspase-3 in a time-dependent manner, inducing apoptotic cell death via a caspase-dependent pathway (Fig. 2C). Furthermore, when AGS-EBV cells were treated with zeocin, the expression of γH2AX was also enhanced along with the increase in the number of eGFP-expressing cells (Supplementary Fig. 1C). These results suggest that both zeocin and bleomycin activate DDR signaling and enhance apoptosis in AGS-EBV and GES1-EBV cells.

### Zeocin induces EBV-positive cell death in a p53-dependent manner

Next, we examined the cytotoxic effects of zeocin in MKN28-EBV and EBV-negative MKN28 cells with mutated p53. The IC_50_ of zeocin on MKN28-EBV cells and MKN28 cells were 1,106 and 1,164 µg/mL, respectively (Fig. 3A). There was no significant difference in cytotoxicity between MKN28-EBV and MKN28 cells (Fig. 3A). Although zeocin treatment induced both p-ATM and γH2AX expression in MKN28-EBV cells, we could not detect the expression of cleaved caspase-3 (Fig. 3B). Consistent with the results obtained from MKN28-EBV cells, knocking-down the wild-type p53 in AGS-EBV cells reduced Zeocin sensitivity (Fig. 3C). These results suggested that zeocin induces apoptosis in EBV-positive cells via a p53-dependent pathway.

**FIG 3.**
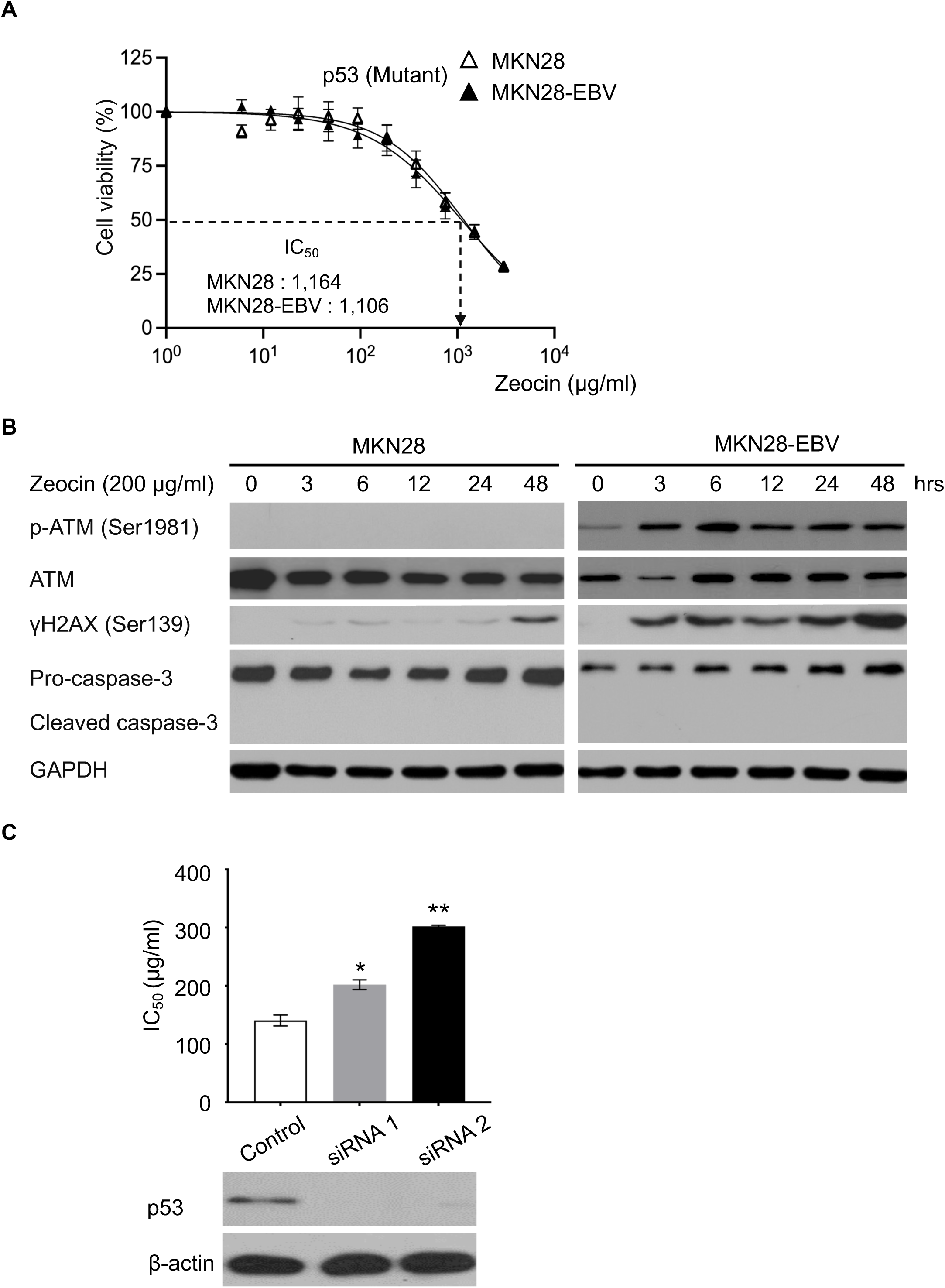
p53 is responsible for EBV-specific DDR enhancement by zeocin. **(A)** Viability of MKN28 and MKN28-EBV cells with p53 mutations after treatment with different concentrations of zeocin. Relative cell viability was calculated by comparison with that of DMSO-treated cells. White triangles, MKN28 cells; black triangles, MKN28-EBV cells. **(B)** Time-dependent expression of DDR-related proteins in MKN28 and MKN28-EBV cells harboring p53 mutations following zeocin treatment. GAPDH was used as an internal control. **(C)** Zeocin-induced death of AGS-EBV cells transfected with p53 siRNA. Upper panel: IC_50_ values for zeocin in siRNA-treated cells. White column: control siRNA; grey column: p53 siRNA 1; black column: p53 siRNA 2. Lower panel: p53 protein expression. β-actin was used as internal control. All data are representative of at least three independent experiments. _*_: *P* <0.05, _**_: *P* <0.01.

### EBV lytic genes, BZLF1 plays an important role in zeocin-induced apoptosis

Next, we examined whether zeocin-induced apoptosis promoted EBV lytic infection and viral production. To measure released EBV from zeocin-treated AGS-eGFP-EBV cells, the culture supernatant was added to the EBV-negative Daudi (-) cell culture (Fig. 4A, left). Viral titers were determined by detecting eGFP-positive cells after eGFP-EBV infection. The culture supernatant obtained from zeocin-treated AGS-eGFP-EBV cells contained approximately 2.5 times more infectious virus than the untreated cells (Fig. 4A, right), with a significance of *P* <0.005. After zeocin treatment, the BZLF1 protein expression increased over time (Fig. 4B). Following zeocin treatment, messenger RNA (mRNA) expression of lytic genes such as BZLF1 (immediate early), BALF5 (early), and BLLF1 (late) increased from 200–900 folds (Fig. 4C). However, when p-53 mutated MKN28-EBV cells were treated with zeocin, BZLF expression was very weak even after 48 h (Supplementary Fig. 2A), changes in BZLF1, BALF5, and BLLF1 mRNAs were within 2-folds and lytic infection was not induced (Supplementary Fig. 2B). To confirm the critical role of BZLF1, AGS-EBV-BZLF1 KO cells were treated with zeocin. The AGS-EBV-BZLF1 KO cells showed higher resistance to zeocin compared to AGS-EBV cells, as evidenced by the IC_50_ of 1,536 µg/ml (Fig. 4D). Compared to AGS cells infected with wild-type EBV, AGS cells infected with EBV-BZLF1 KO exhibited a significant decrease in apoptosis, from 35.71–14.94% (Fig. 4E). Although zeocin treatment activated the DDR signal (p-ATM and γH2AX) in both AGS-EBV and AGS-EBV-BZLF1 KO cells, AGS-EBV-BZLF1 KO cells did not show the increase in cleaved caspase-3 expression (Fig. 4F). Furthermore, changes in the expression of early and late genes were not observed in AGS-EBV BZLF1 KO cells treated with zeocin (Supplementary Fig. 3A). Zeocin treatment increased the viral genome copy number by >100-fold in AGS-EBV cells, but not in AGS-EBV BZLF1 KO cells (Supplementary Fig. 3B and 3C). These results indicated that BZLF1 plays a critical role in zeocin-induced apoptosis, cell death, and lytic induction through the DDR pathway.

**FIG 4.**
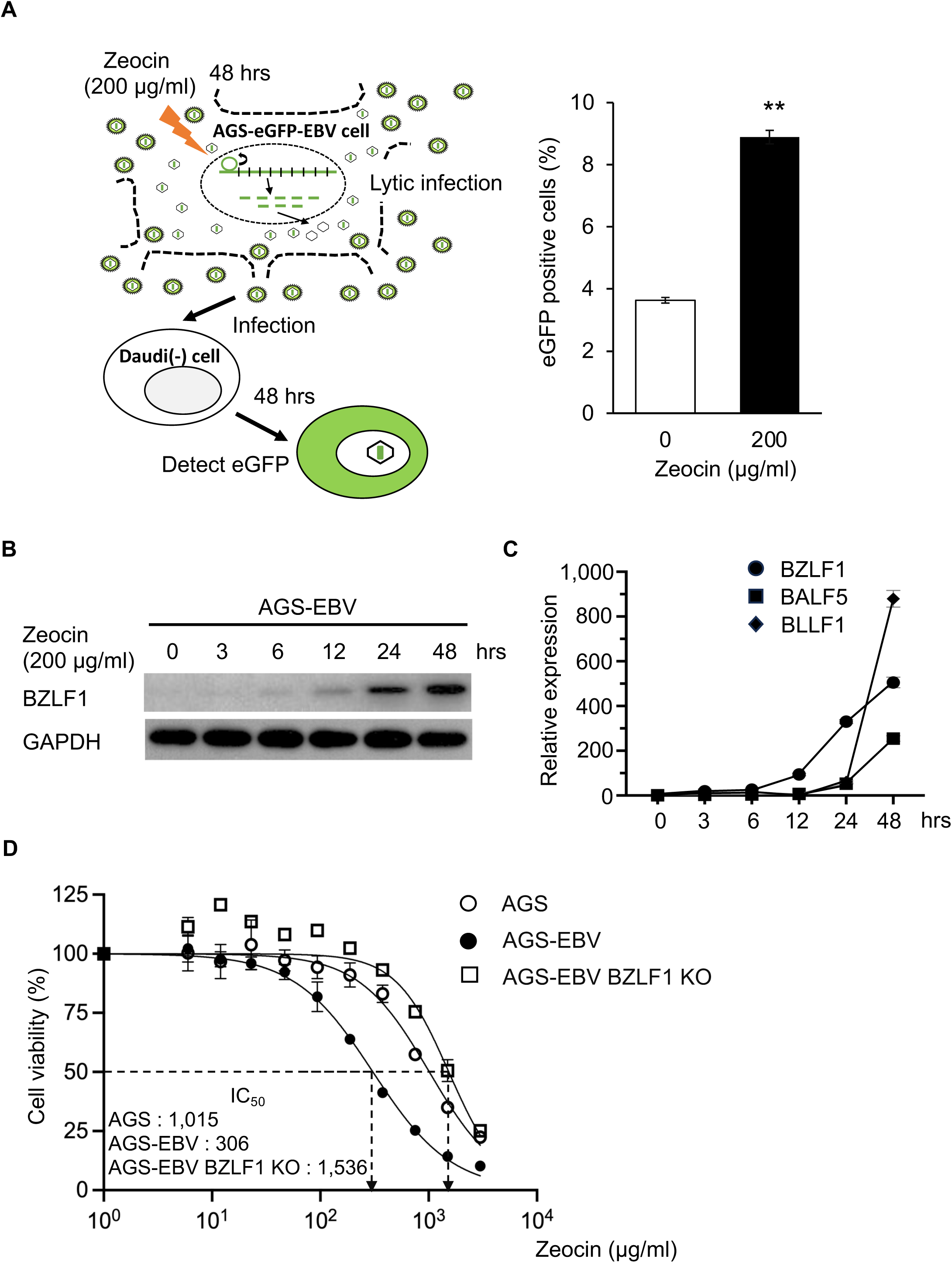
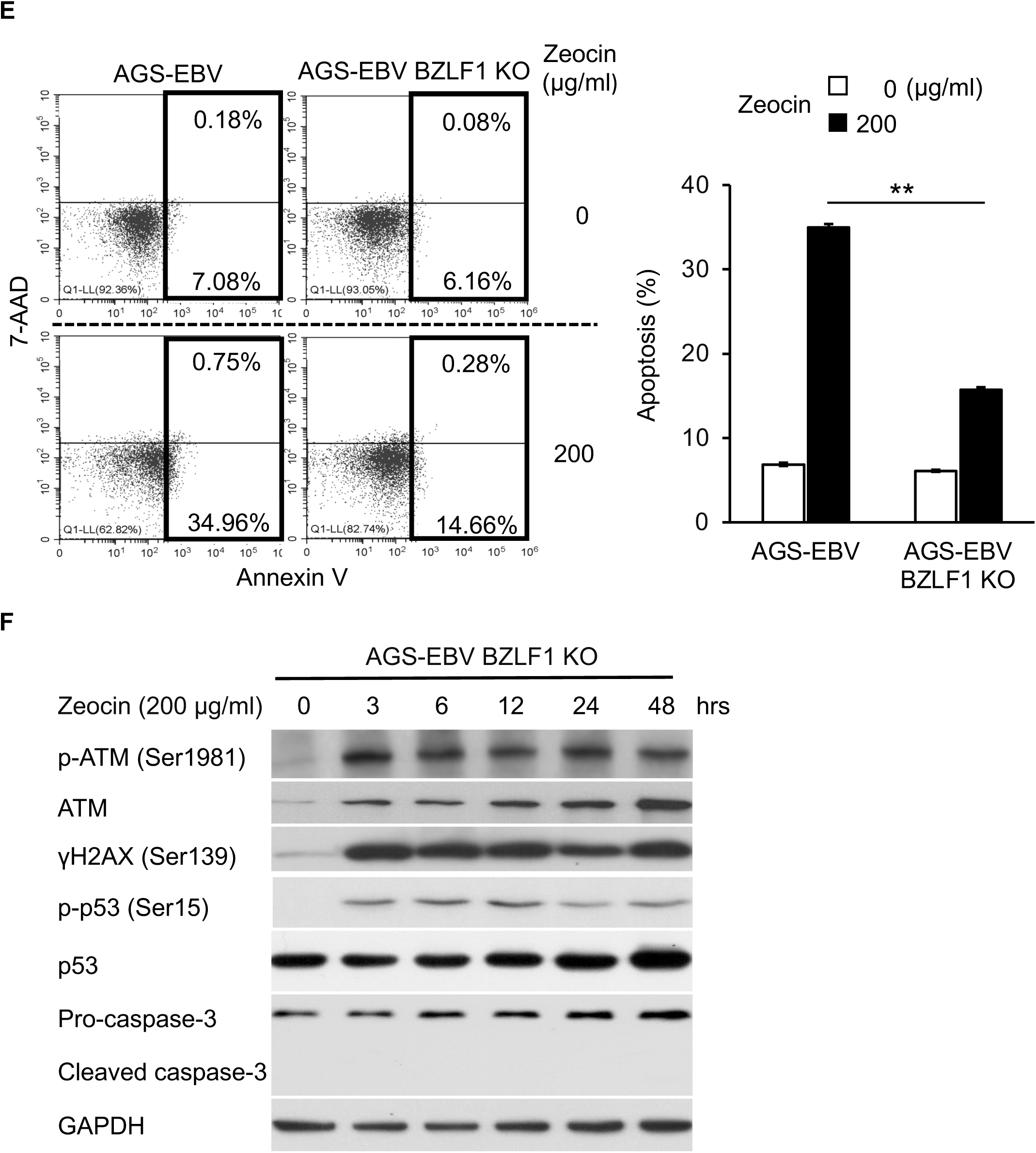
Zeocin-induced cell death in EBV-positive cells leads to subsequent lytic viral replication. **(A)** Detection of infectious virus from zeocin-treated EBV-positive epithelial cell culture supernatant. Left: Schematic presentation to measuring infectious virus. Right: Detection of eGFP-positive cells in Daudi (-) cells infected with supernatant eGFP-EBV. **(B)** Expression of BZLF1 in AGS-EBV cells after zeocin treatment. GAPDH was used as internal control. **(C)** Expression of lytic EBV genes in AGS-EBV cells after zeocin treatment. Black circle: BZLF1 (immediate early), Black square: BAL5 (early), Black diamond: BLLF1 (late). The mRNA expression was normalized by GAPDH. The level of mRNA expression in DMSO-treated cells was set to 1. **(D)** Cell viability in AGS cells after zeocin treatment. The relative viability of zeocin treated cells was calculated by comparing with DMSO treated cells. White circle: AGS cells, Black circle: AGS-EBV cells, White square: AGS-EBV BZLF1 KO cells. **(E)** Apoptosis after zeocin treatment in AGS-EBV cells and AGS-EBV BZLF1 KO cells. Left: FACS analysis, Early apoptosis: 7-AAD^-^/Annexin V^+^, Late apoptosis: 7-AAD^+^/Annexin V^+^. Right: Frequency of apoptosis after zeocin treatment. Apoptosis was calculated by early plus late apoptosis. **(F)** Time-dependent expression of DDR-related proteins in AGS-EBV BZLF1 KO cells after zeocin treatment. GAPDH was used as internal control. All experiments are representative of at least three independent experiments. _*_: *P* <0.05, _**_: *P* <0.01.

### LMP2A enhances DDR related lytic infection

As shown in Fig. 4F, zeocin treatment of AGS-EBV BZLF1 KO cells did not show caspase-3 activation, but induced phosphorylation of ATM and γH2AX. We assumed that any of the EBV latent genes expressed in AGS-EBV BZLF1 KO cells induced DDR. Among viral latent genes, LMP2A was assumed to induce DDR. We introduced the LMP2A-expressing plasmid into virus-free AGS cells, treated the cells with zeocin, and examined the activation of DDR. In LMP2A transfected cells, γH2AX expression was upregulated in the presence of zeocin (Fig. 5A). AGS-EBV LMP2A KO cells exhibited an IC_50_ of 1,076 µg/mL, which was three times higher than AGS-EBV cells (Fig. 5B). Moreover, γH2AX expression was less weakly induced in zeocin treated AGS-EBV LMP2A KO cells compared with AGS-EBV cells (Fig. 5C). In addition, caspase-3 activation was not detected in zeocin-treated AGS-EBV LMP2A KO cells (Fig. 5C). After zeocin treatment, the expression of lytic genes increased over time in wild-type virus-infected AGS-EBV cells, but the enhanced mRNA expression of BZLF1, BALF5, and BLLF1 decreased from 1/30 to 1/100 in AGS-EBV LMP2A KO cells compared to that in AGS-EBV cells (Fig. 5D). Additionally, AGS-EBV LMP2A KO cells reduced the number of EBV copies by less than one-fifth compared to AGS-EBV cells (Fig. 5E). These results show that LMP2A enhances DDR signaling and induces lytic infection in AGS-EBV cells.

**FIG 5.**
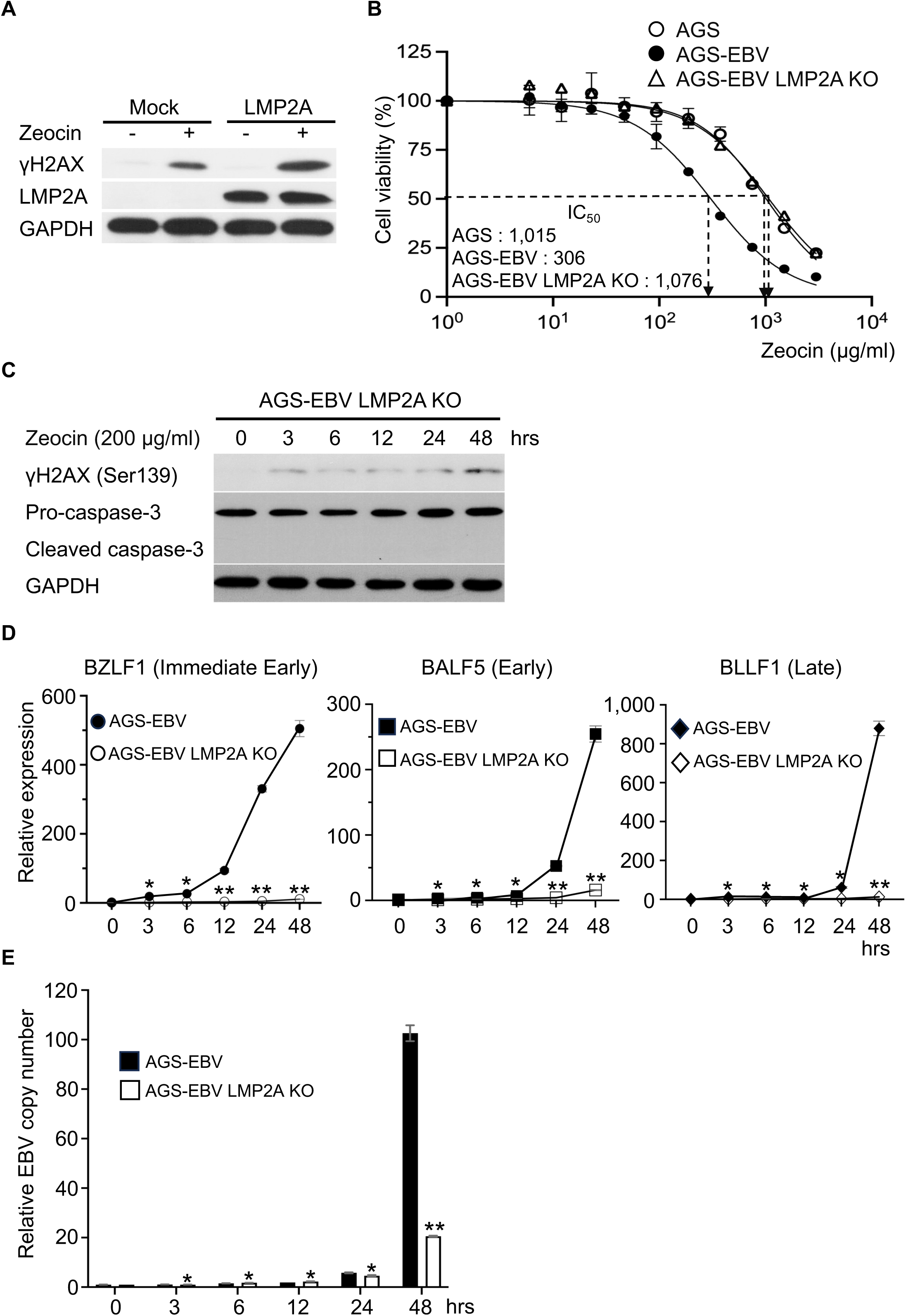
Viral LMP2A enhances zeocin-induced DDR. **(A)** Zeocin-induced DDR in AGS cells and LMP2A-expressing AGS cells. GAPDH was used as internal control. **(B)** Cell viability in AGS cells, AGS-EBV cells, and AGS-EBV LMP2A KO cells after zeocin treatment. The relative viability of zeocin treated cells was calculated by comparing with DMSO treated cells. White circle: AGS cells, Black circle: AGS-EBV cells, White triangle: AGS-EBV LMP2A KO cells. **(C)** Expression of DDR-related proteins in AGS-EBV LMP2A KO cells after zeocin treatment. GAPDH was used as internal control. **(D)** Expression of lytic EBV genes in AGS-EBV LMP2A KO cells after zeocin treatment. BZLF1 (immediate early), BAL5 (Early), and BLLF1 (Late) were expressed by circle, square, and diamond, respectively. Black circle, square, and diamond: AGS-EBV cells, White circle, square, and diamond: AGS-EBV LMP2A KO cells. The relative mRNA expression was normalized by GAPDH. The level of mRNA expression in DMSO-treated cells was set to 1. **(E)** Relative viral copy number in AGS-EBV and AGS-EBV LMP2A KO cells treated with zeocin. Black square: AGS-EBV cells, White square: AGS-EBV LMP2A KO cells. Viral copy number was calculated in AGS-EBV and AGS-EBV LMP2A KO cells, respectively, by setting GAPDH gene copy number of DMSO-treated cells to 1. All experiments are representative of at least three independent experiments. _*_: *P* <0.05, _**_: *P* <0.01.

## DISCUSSION

EBV reactivates from latent to lytic infections as part of its life cycle. When BZLF1, the master switch for reactivation, is activated, the EBV genome is amplified >100 times (4). During EBV replication, p53 binds directly to and activates the BZLF1 promoter (11). Therefore, when p53 expression is suppressed in EBV-infected cells, the BZLF1 promoter is not activated by chemicals that induce lytic infection (Chang et al., 2008). In addition, p53 expression is often normal in EBV-positive epithelial tumors, such as gastric cancer and nasopharyngeal carcinoma (14, 37). Therefore, when DDR occurs in EBV-positive epithelial tumors, p53 is activated, inducing EBV lytic infection, and the infected cells undergo apoptosis. Detailing the molecular mechanisms underlying these reactions will aid in the development of drugs specific to EBV-positive epithelial tumors.

In addition, induction of EBV lytic infection by BZLF1 activation induces the expression of viral regulatory factors such as BRLF1 and BRRF1, forming a positive feedback loop and causing DDR (38). Additionally, the BGLF4 kinase, which is expressed during the viral lytic infection phase, phosphorylates Tat-interactive protein, 60 kDa and ATM, resulting in DDR (39, 40). In contrast, the viral latent genes Epstein-Barr nuclear antigen 1, EBNA3, and latent membrane protein 1 suppress the DDR (41–44). EBV miRNAs also cooperatively downregulate ATM (45). Therefore, apart from overexpression (46), latent EBV genes are believed to suppress DDR.

Regarding LMP2A, little research has been conducted on its relationship with DDR. Wasil et al., demonstrated that DDR was enhanced when HEK293 cells were overexpressed with the LMP2A gene and treated with the anticancer drug etoposide (47). However, this observation raised the possibility that the heightened DDR response was due to the overexpression of LMP2A beyond normal physiological levels. In contrast, our study illustrated that gastric epithelial cells infected with recombinant EBV lacking LMP2A exhibited reduced DDR and cytotoxic response induced by zeocin treatment compared to cells infected with wild-type EBV (Fig. 5A). These findings strongly suggested that LMP2A plays a role in promoting drug-induced DDR.

LMP2A mimics pre-BCR signal transmission and plays a pivotal role in signaling in EBV-infected cells (21). The pre-BCR is responsible for transmitting proliferation signals and initiating DDR, which subsequently triggers the reconfiguration of immunoglobulin genes (48). Despite this, B cells possess an inherent mechanism for DDR suppression via interleukin-7 receptor signaling (48), effectively mitigating excessive DDR. EBV primarily infects B cells and exhibits a lower frequency of infection in epithelial cells (49, 50), implying that its evolutionary adaptation was not primarily centered on infecting epithelial cells. Consequently, it is plausible that EBV-positive epithelial cells lack certain mechanisms, such as interleukin-7 receptors, to suppress heightened DDR.

Moreover, because EBV-positive gastric epithelial cells show type I latent infection and are notably deficient in EBNA3 or latent membrane protein 1, which typically suppress DDR (20), these cells frequently display intensified DDR. This heightened vulnerability to DDR could potentially facilitate increased production of viral progeny through the induction of lytic infection. Studies have indicated higher numbers of EBV DNA copies in the blood of patients with EBV-positive epithelial cancer than in healthy individuals (51, 52). This is likely because frequent EBV reactivation is observed in EBV-positive epithelial cells (53).

In this study, we showed that LMP2A enhanced the DDR and induced p53-dependent apoptosis in EBV-positive gastric epithelial cells by facilitating lytic infection (Fig. 6). The use of drugs as anticancer agents that induce cell death in EBV-associated cancers has also been discussed. The use of ganciclovir or HDAC inhibitors in combination with drugs that induce EBV lytic infection has been reported (54, 55). Another report recommended the use of poly ADP-ribose polymerase inhibitors because EBV infection activates the STAT3 pathway and BRCAness (56). Bleomycin has been used to treat Hodgkin’s lymphoma (57, 58). Thus, we expect to develop specific therapies for EBV-associated cancers using drugs that induce DDR, such as the bleomycin family. Since DDR responsiveness occurs even more strongly in EBV-positive epithelial cancers that express LMP2A, we believe that bleomycin family drugs have a high potential for application in oncolytic therapy for EBVaGC.

**FIG 6.**
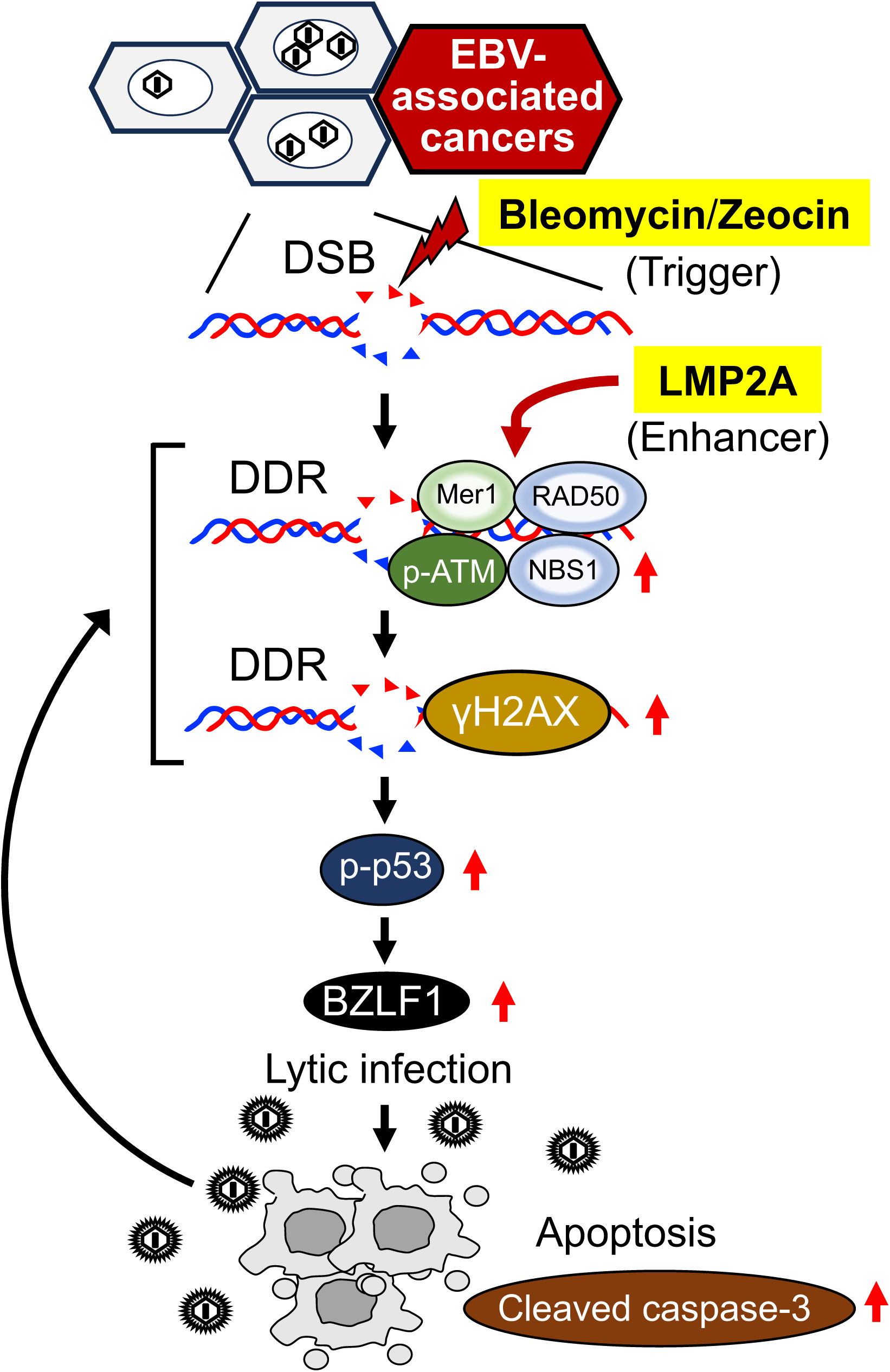
Graphic summary of the current work. DDR induced by the bleomycin family is enhanced by viral latent gene LMP2A. DDR is activated by p53 and LMP2A and leads to lytic EBV infection. Viral lytic infection further instigates DDR and induces apoptosis. DSB: double-stranded DNA breaks.

## ABBREVIATIONS

ATM: ataxia-telangiectasia mutated
BZLF1: *Bam*HI Z fragment leftward open reading frame 1
DDR: DNA damage response
EBV: Epstein-Barr virus
EBVaGC: EBV-associated gastric cancer
EBERs: EBV-encoded small RNAs
eGFP: enhanced green fluorescence protein
GAPDH: glyceraldehyde triphosphate dehydrogenase
γH2AX: histone H2AX phosphorylated on Ser 193
LMP2A: latent membrane protein 2A
siRNA: small RNA interference

## ACKNOWLEDGMENTS

This work was supported by JSPS KEKNHI (grant number 22K07101 and 21K07054), the Otsuka Toshimi Scholarship foundation (grant number 20-112 and 21-62), the SGH Scholarship Foundation and JST SPRING (grant number JPMJSP2155). The authors thank Noriko Ito, Aoi Osakada, Yukiko Sakamoto, Hayato Odaka, Momoko Fukuda and Emi Kurauchi for their assistance.

---

**Supplementary Fig. 1.**
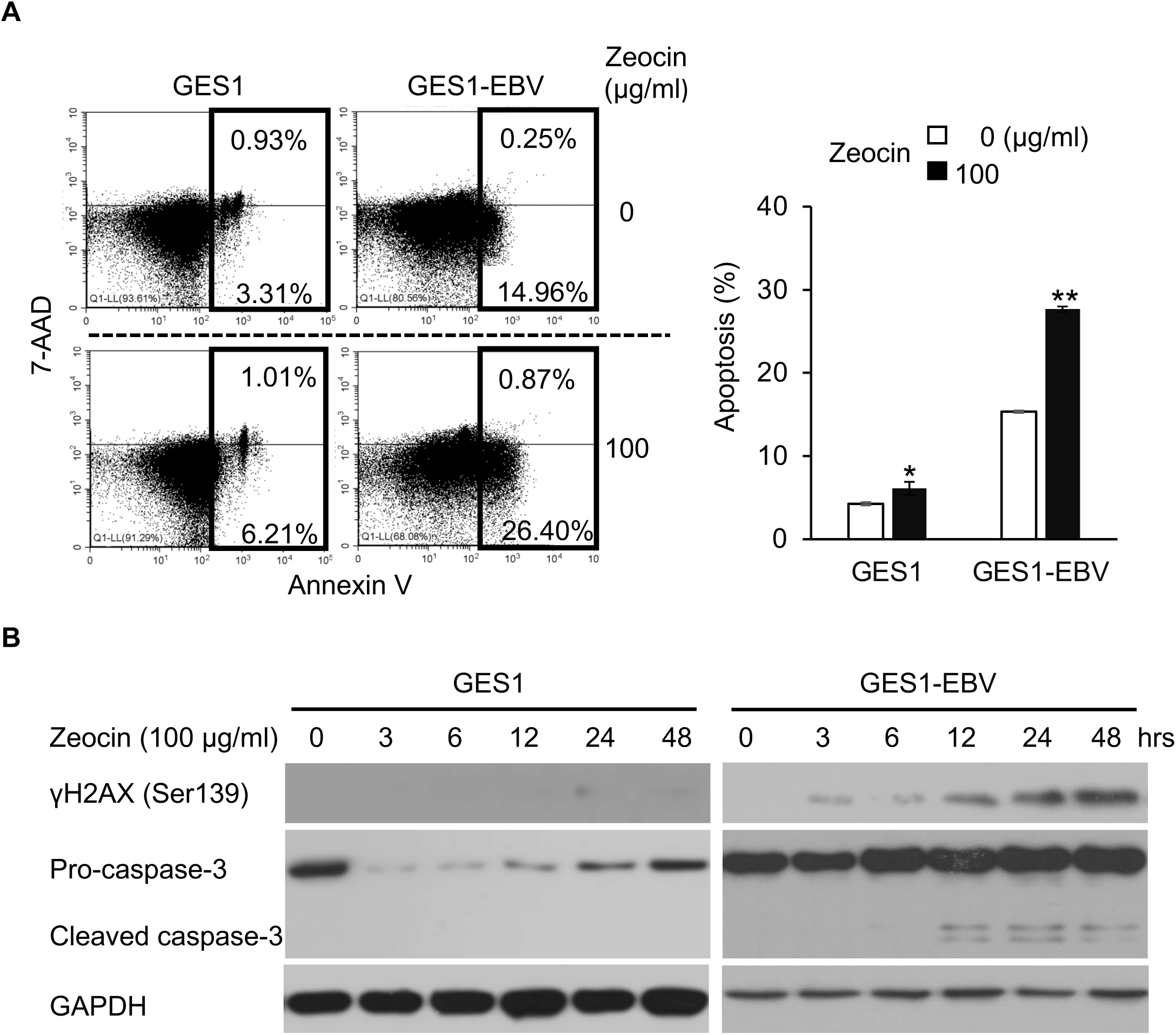
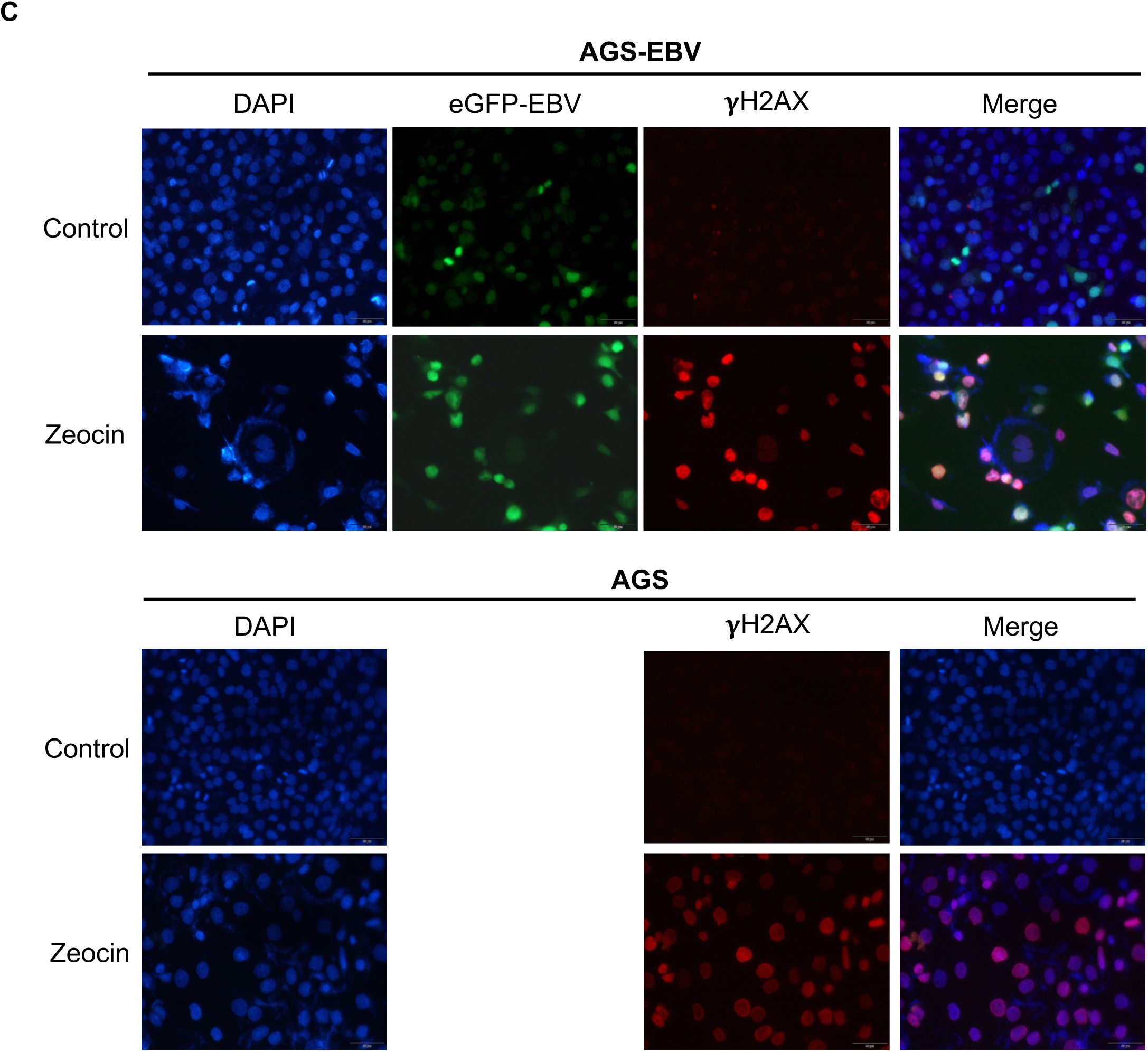

**Supplementary Fig. 2.**
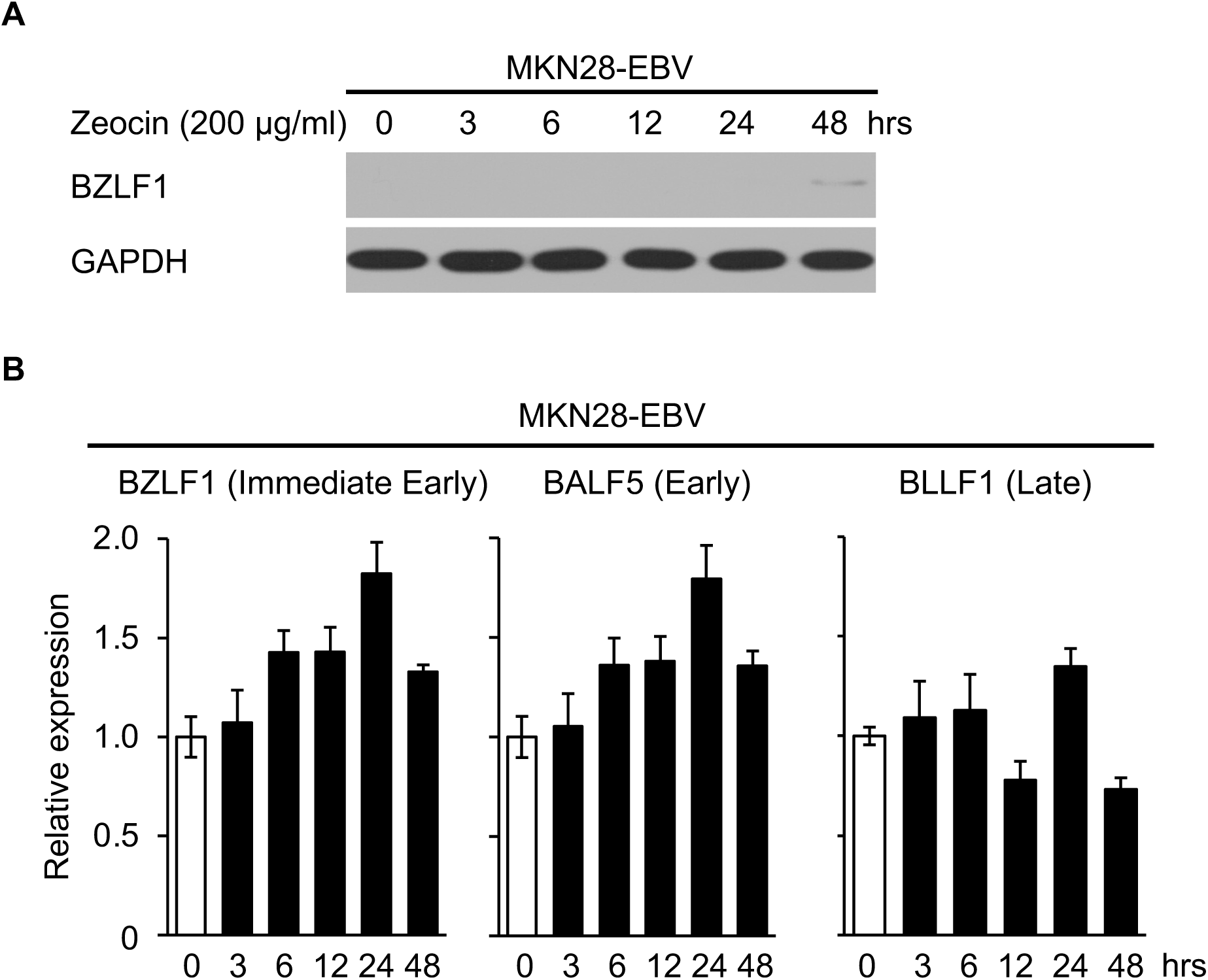

**Supplementary Fig. 3.**
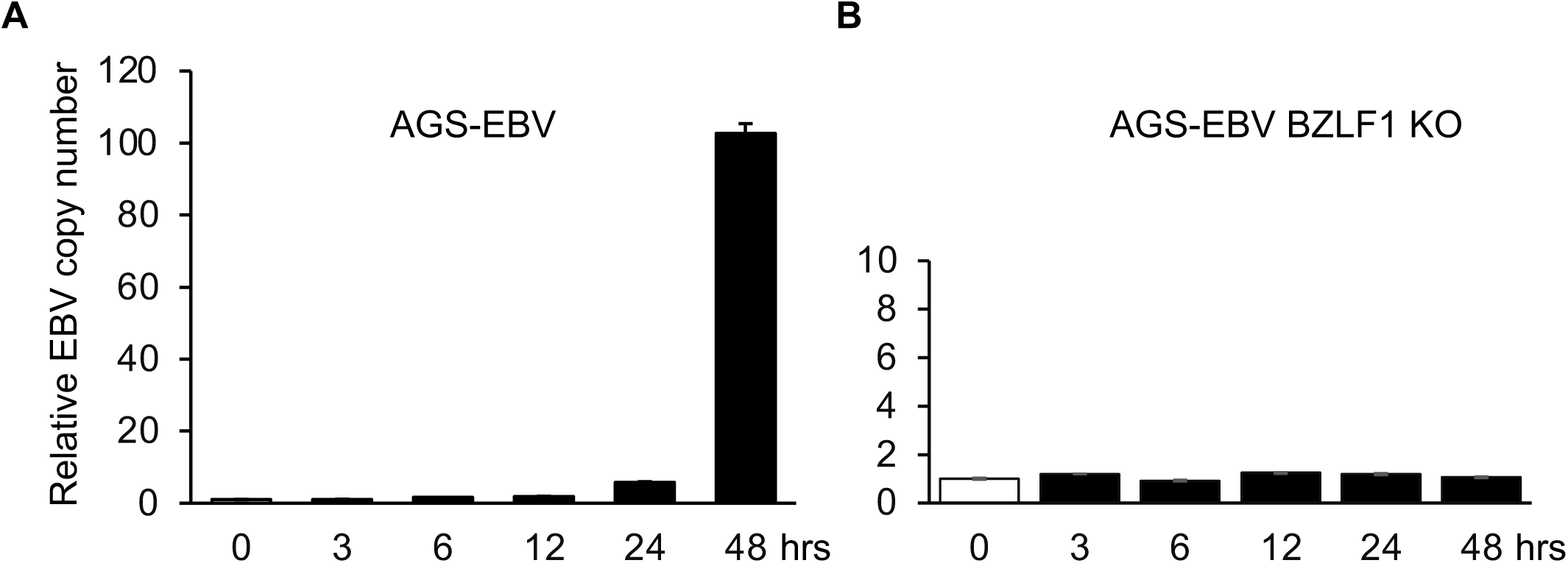

